# IMPACT: a web server for exploring immunotherapeutic predictive and cancer prognostic biomarkers

**DOI:** 10.1101/2023.01.31.526401

**Authors:** Yutao Liu, Yundi Zhang, Wenchuan Xie, Jing Zhao, Yiting Dong, Chunwei Xu, Yanan Wang, Man Li, Guoqiang Wang, Xin Zhu, Wenxian Wang, Kequan Lin, Huafei Lu, Yusheng Han, Leo Li, Jianchun Duan, Shangli Cai, Jie Wang, Zhijie Wang

## Abstract

Immune checkpoint inhibitors (ICIs) are a breakthrough in oncology treatment, and studies of screening predictive biomarkers of ICIs are emerging. We developed a web server named IMPACT (http://impact.brbiotech.com/) to thoroughly explore immunotherapeutic predictive or prognostic biomarkers. IMPACT contains a large dataset of 6,276 patients treated with ICIs and integrates 11 well-designed function modules, enabling an in-depth solution for biomarkers exploration. Compared with the existing tools, IMPACT was implemented with one exclusive module for interaction analysis and several optimized conventional functions for discovering novel biomarkers. Specifically, the interaction analysis of biomarker-treatment effect is essential to determine whether a biomarker is predictive and/or prognostic for ICIs. Moreover, several optimized functions allow complicated biomarker exploration, including customized selections of variant types in more detail, automatically screening meaningful co-mutations among multiple genes, and selecting cut-off values for gene expression biomarkers. In summary, IMPACT is a comprehensive analysis resource to facilitate biomarker research of ICIs.

## Introduction

Immune checkpoint inhibitors (ICIs), one of the most important breakthroughs in oncology, have dramatically improved patients’ prognoses in a range of tumors ^1^. However, only a subset of patients can benefit from ICIs ^2^. Biomarkers that can effectively distinguish immune responders from non-responders are crucial to improve the clinical use of ICIs. Currently, several biomarkers for ICIs have been identified, including PD-L1 expression ^3^, tumor mutational burden (TMB) ^3^, *LRP1B* mutation ^4^, *STK11* mutation ^5,6^, *NOTCH* mutation ^7^, *KEAP1* mutation ^5^, and expression signatures of tumor infiltration lymphocytes ^3^. However, the relationship between various biomarkers is intricate and up to now there is no omnipotent biomarker suitable for all cancer types and clinical scenarios. Therefore, urgently needed are novel or combined predictive biomarkers for ICIs. Currently, some tools have been developed based on public datasets and are available for exploring potential biomarkers, such as CAMOIP ^8^, Cancer-Immu ^9^, CRI-iAtlas ^10^, and the cBioPortal ^11^. However, one analysis that is critical to biomarker exploration has been overlooked: interaction analysis between a biomarker and treatment that is indispensable for determining whether the biomarker is an ICIs-specific predictor or a treatment-independent cancer prognostic factor ^12^. In addition, several conventional functions have excellent optimized space in those existing tools, including co-mutation exploration, self-defined biomarkers based on specific variant types, and optimal cut-off value selection for gene expression biomarkers. Thus, we developed a web server, IMPACT (<u>IM</u>munotherapeutic <u>P</u>redictive <u>A</u>nd <u>C</u>ancer prognos<u>T</u>ic biomarkers), compatible with both public and in-house genomic and transcriptomic data. In summary, IMPACT is a publicly available and user-friendly platform for exploring immunotherapeutic predictive and cancer prognostic biomarkers based on genomic or transcriptome data, which can facilitate the investigation and visualization of predictive biomarkers particularly for immunotherapy, relevant interaction effects, and biological mechanisms.

## Methods

### Data collection

The genomic or transcriptomic data of patients treated with ICIs were collected from 24 public datasets and 3 in-house datasets, corresponding to 6,276 samples across 10 cancer types (**Supplementary Table S1**). In addition, 1 in-house non-ICI dataset and 48 public non-ICI datasets were curated from databases of TCGA, GEO, CPTAC, ICGC, and OncoSG. In detail, the genomic data were generated based on the whole exome sequencing (WES) or the next-generation sequencing (NGS) panel, while the transcriptomic data were derived from the whole transcriptome sequencing (WTS), microarray, or specific RNA panel. The TMB score was obtained from the original datasets if available; otherwise, the TMB score of a sample was defined by its total number of non-synonymous mutations. The information on objective response and progress-free survival (PFS) was obtained from these original studies, of which 25 were evaluated by the Response Evaluation Criteria In Solid Tumors (RECIST) v1.1, 1 was by the immune-related RECIST (irRECIST), and 1 was self-defined. Overall survival (OS) was also obtained from these original studies. The OS of three in-house datasets was defined as the time from the treatment date to the date of death or the end of follow-up.

### Data processing

All clinical and molecular data were converted into a uniform format. For transcriptomic data, the fragmentation per kilobase million (FPKM) was converted to the transcript per million bases (TPM), and then all the TPM was standardized with log_2_(TMP + 0.001). Read counts were converted into log_2_(CPM + 0.001). Signatures of immune and oncogenic pathways were obtained from a previously published literature ^13^ or MSigDB Database (https://www.gsea-msigdb.org/gsea/msigdb). Then the ssGSEA algorithm was applied to calculate the enrichment scores of these signatures.

### Statistics

Kaplan-Meier (KM) survival curves and univariable/multivariable Cox regression models were used to analyze the associations between biomarkers and OS/PFS. A random-effects meta-analysis model was used to pool the results from multiple studies with the “meta” and “survminer” R packages. The ggforest function in the “survminer” package was modified for forest plotting. Continuous variables between two groups were compared using the Mann-Whitney U test and categorical variables using the Chi-square test or Fisher exact test. P value for multiple tests was adjusted by the Benjamini-Hochberg in the correlation analysis module between biomarkers and gene expression.

### Uploading function

IMPACT allows users to upload their own datasets. Uploaded data should include clinical outcome information and their related genetic and/or transcriptomic data. For somatic mutation, each patient should be clearly annotated with mutation (MUT) or wild type (WT). Transcriptomic data should be normalized and transformed with log2. An example of the data format is provided on the “User-defined” page of IMPACT.

### Parameter customization

The key parameters can be customized during analysis by users to fulfill their specific needs, such as variant-type defining mutation (MUT) and wild type (WT), cut-off values for continuous variable categorization, graph colors, and legend positions. These customization can greatly increase the utility and flexibility of IMPACT and help users generate desired figures directly from IMPACT.

### Data availability statement

All the data used in IMPACT can be downloaded from our server at http://impact.brbiotech.com (http://www.brimpact.cn/).

### Code availability statement

All the code can be accessed at https://github.com/wenchuanxie/IMPACT.

## Results

### Overview of IMPACT

IMPACT was developed for systematically investigating predictive or prognostic biomarkers, relevant interaction effects, and biological mechanisms (**Figure 1**), therefore contributing to untangling the current clinical challenges, such as (i) understanding ICIs biomarkers with synergistic or interaction effects, (ii) identifying a predictive biomarker specified for ICIs, and (iii) exploring the functional mechanism of the biomarkers of interest. Eleven function modules were implemented in IMPACT, including PredExplore (Immunotherapy Predictive Biomarker Exploration), ProgExplore (Cancer Prognostic Biomarker Exploration), Survival Analysis (composed of Kaplan-Meier Curve, Cox Regression, Subgroup analysis, Cutpiont Analysis, Immunotherapy Response), Interaction Analysis, Immunogenicity Analysis, Microenvironment Analysis, and Mutation Profiles.

**Figure 1.**
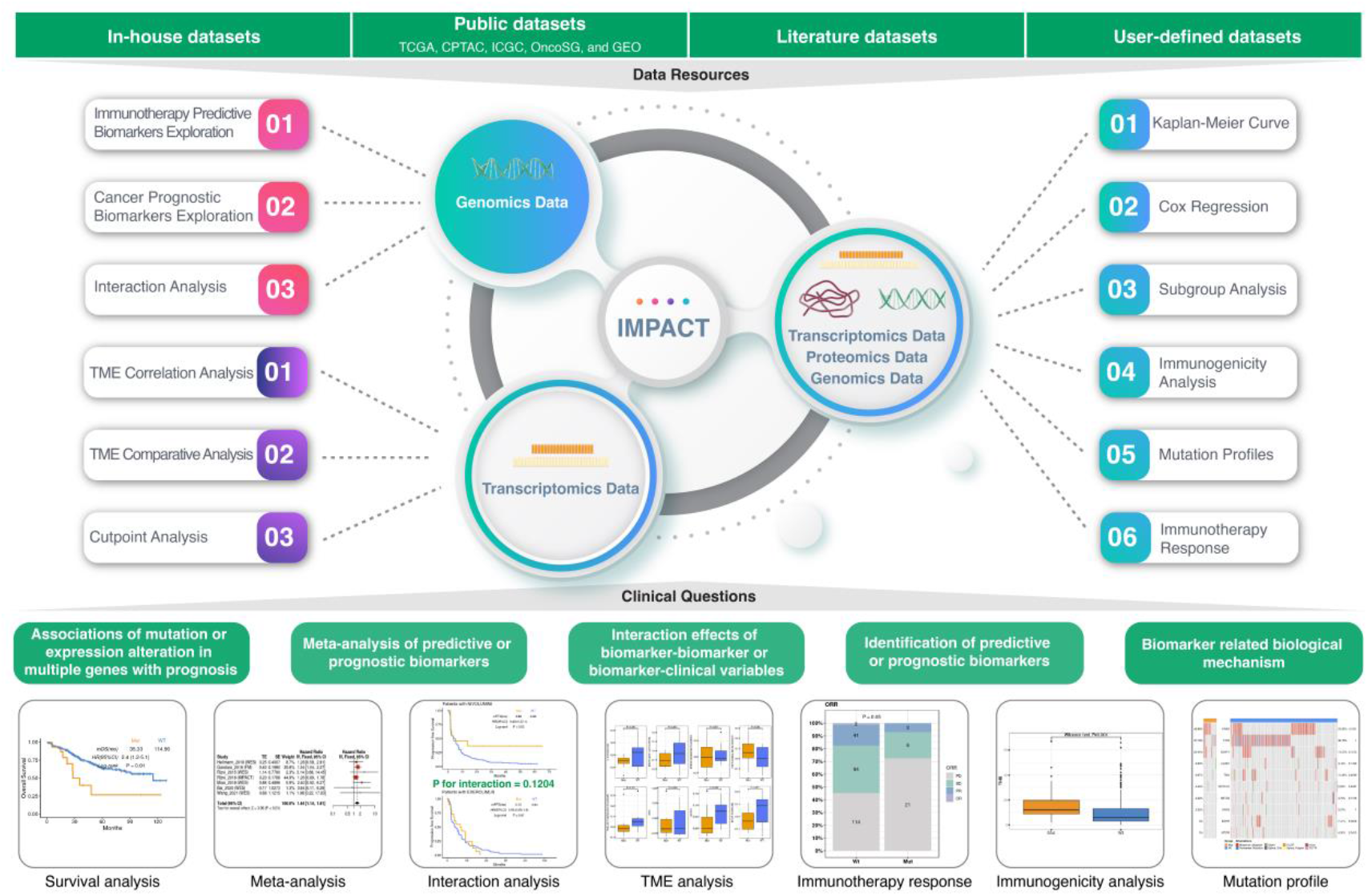
The overview of IMPACT

We conducted literature review and found five existing tools similar to IMPACT. A detailed comparison among them in terms of data sources, clinical outcomes, and functions were summarized in **Table 1**. Exclusively, IMPACT integrates an “Interaction Analysis” module that can identify interaction effects between biomarkers to distinguish predictive and prognostic biomarkers. As for some common features shared among these tools, we improved their utility and capability in IMPACT. For instance, (i) Co-mutation biomarker screening. Co-existing genomic alterations in oncogenic drivers could represent certain categories of molecular diversity and interplay that influences tumor development and therapeutic outcomes. To discover potential biomarkers of co-mutations, IMPACT can automatically analyze all the pair-wise combinations of mutations up to 50 selected genes to explore potential co-mutation biomarker (see the example in **Supplementary Table S2**); (ii) Mutation type selection. Since different types of mutations have varied influence on gene functions, e.g, a truncating mutation of a gene (nonsense, frame-shift insertion/deletion, and splice) may result in a loss of function, while a missense mutation of the same gene may reduce or increase the function, it is necessary to take mutation types into consideration when defining the mutation or wild-type. IMPACT allows users to specifically select the variant type of mutation (see the example in **Supplementary Figure S1A-B**); (iii) Cut-off value optimization. The optimal cut-off values for continuous biomarkers, such as gene expression and TMB, are critical to stratify patients and to explore the effects of cut-off value selection on the associations between biomarkers and survival. We also provided cut-off value analysis in IMPACT (see the example in **Supplementary Figure S1C-D**); (iv) Customized confounder analysis. To determine whether a biomarker is an independent predictive or prognostic biomarker, IMPACT allows users to customize the multivariable analysis by selecting other confounders, including clinical factors of interest or related to prognosis; (v) Comparison between responders and non-responders. IMPACT can compare differentially mutated/expressed genes or clinical factors between response and non-response groups to identify significant factors associated with the response to ICIs (see the example in **Supplementary Figure S1E-J**). Here, two case studies are provided to show how IMPACT may facilitate researchers to comprehensively explore biomarkers.

**Table 1.**
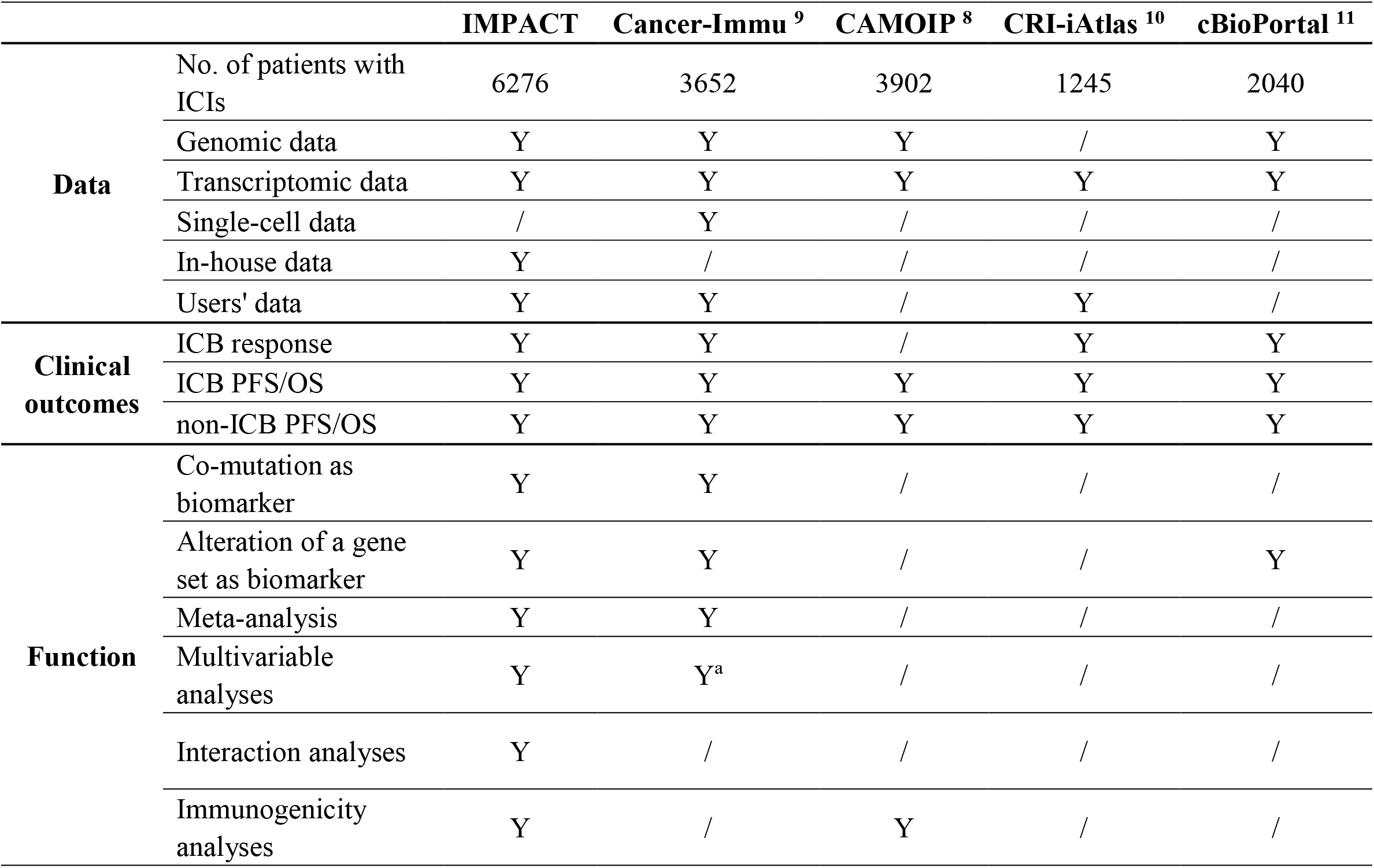
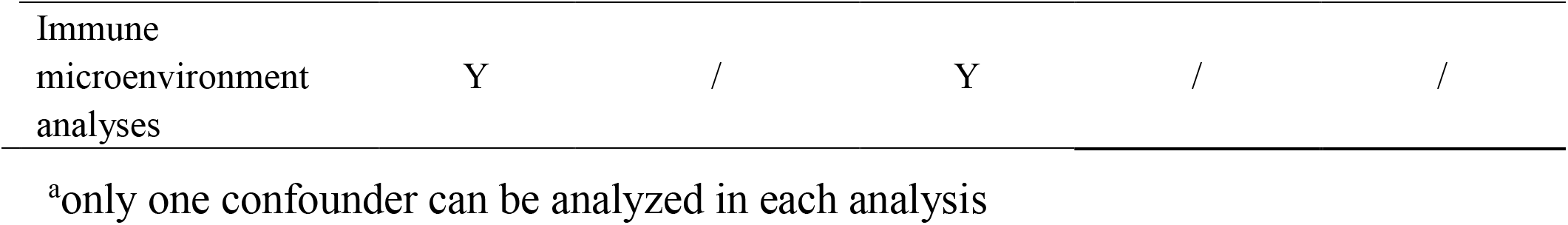
Comparison between IMPACT and other existing tools

|  |  | IMPACT | Cancer-Immu<sup>9</sup> | CAMOIP<sup>8</sup> | CRI-iAtlas<sup>10</sup> | cBioPortal<sup>11</sup> |
| --- | --- | --- | --- | --- | --- | --- |
| **Data** | No. of patients with ICIs | 6276 | 3652 | 3902 | 1245 | 2040 |
|  | Genomic data | Y | Y | Y | / | Y |
|  | Transcriptomic data | Y | Y | Y | Y | Y |
|  | Single-cell data | / | Y | / | / | / |
|  | In-house data | Y | / | / | / | / |
|  | Users' data | Y | Y | / | Y | / |
| **Clinical outcomes** | ICB response | Y | Y | / | Y | Y |
|  | ICB PFS/OS | Y | Y | Y | Y | Y |
|  | non-ICB PFS/OS | Y | Y | Y | Y | Y |
| **Function** | Co-mutation as biomarker | Y | Y | / | / | / |
|  | Alteration of a gene set as biomarker | Y | Y | / | / | Y |
|  | Meta-analysis | Y | Y | / | / | / |
|  | Multivariable analyses | Y | Y<sup>a</sup> | / | / | / |
|  | Interaction analyses | Y | / | / | / | / |
|  | Immunogenicity analyses | Y | / | Y | / | / |

|  |  |  |  |  |  |
| --- | --- | --- | --- | --- | --- |
| Immune<br>microenvironment<br>analyses | Y | / | Y | / | / |

<sup>a</sup>only one confounder can be analyzed in each analysis

### Application 1: To comprehensively explore the associations of *STK11* with prognosis or immune signatures in lung adenocarcinoma in ICIs and non-ICIs datasets

#### 1. Immunotherapy Biomarker and Survival Exploration

Using the “PredExplore” module, we analyzed the association between *STK11* and survival in all available ICIs cohorts of lung adenocarcinoma (LUAD). The results showed that their association was significant in 3 of 4 datasets for OS and 1 of 7 datasets for PFS (all the P values < 0.05, **Supplementary Table S3**). In the meta-analysis, the association was significant for both OS (hazard ratio [HR] = 1.66, 95% CI = 1.29−2.14, **Figure 2A**) and PFS (HR = 1.44, 95% CI = 1.14−1.81, **Supplementary Figure S2A**). Furthermore, Kaplan-Meier curves and clinical response analyses for each dataset can be realized through their corresponding modules under the “Survival Analysis”. Taking the Gandara_2018 dataset as an example, patients carrying the *STK11* mutation (defined as nonsynonymous mutation) had shorter OS and PFS (mOS: 7.33 mo vs 15.61 mo, Log-rank P = 0.005, **Figure 2B**; mPFS: 1.41 mo vs 2.79 mo, Log-rank P = 0.03, **Supplementary Figure S2B**) than those with wild-type *STK11* when treated with ICIs, as well as a higher percentage of progressive diseases (P = 0.05, **Figure 2C**). Furthermore, the result of “Cox regression” demonstrated that the association of the *STK11* mutation and OS remained significant in the multivariable Cox regression analysis (HR = 1.66, 95% CI, 1.08−2.55, P = 0.02, **Figure 2D**). As for PFS, the result of univariable Cox regression was significant, but not in multivariable Cox regression (**Supplementary Figure S2C**). Altogether, these results suggest that the *STK11* mutation could be an independent predictive and/or prognostic biomarker for ICIs while more analyses were needed.

**Figure 2.**
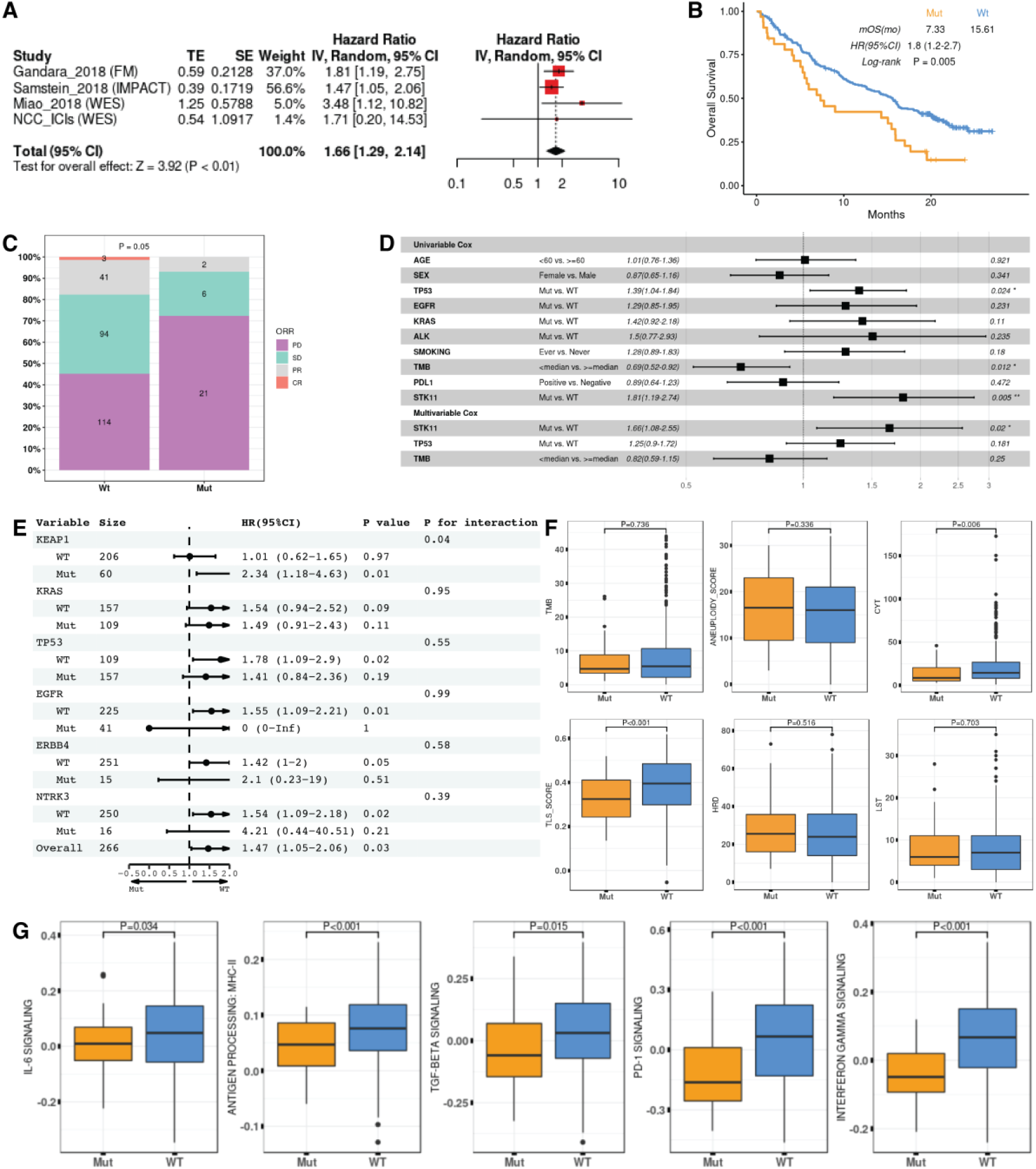
The exploration of *STK11* mutation as a predictive or prognostic biomarker. **(A)** Meta-analysis of *STK11* mutation and OS across LUAD ICIs cohorts. **(B)** The Kaplan-Meier curve of *STK11* mutation and OS on the Gandara_2018 dataset. **(C)** The correlation of *STK11* mutation and objective response rate on the Gandara_2018 dataset. **(D)** The univariable and multivariable cox regression analyses of *STK11* mutation with OS on the Gandara_2018 dataset. **(E)** The interaction analysis between mutation of *STK11* and other genes on the Samstein_2018 dataset. **(F)** and **(G)** The correlation of *STK11* mutation with immunogenicity and immune signatures on the TCGA-LUAD dataset.

#### 2. Subgroup and Interaction Analysis

To further explore the association of the *STK11* mutation and survival in LUAD, subgroup analyses were performed for different stratifications of clinical characteristics using the “Subgroup Analyses” module. The results showed that the association of the *STK11* mutation with OS was consistent in each subgroup (all the P interaction > 0.05, **Supplementary Figure S2D**).

Using IMPACT, users can also investigate the interaction effects between multiple genes. Here, we used the “Mutational Profiles” module to analyze the association of the *STK11* mutation with other gene mutations, and unveiled that the *STK11* mutation was positively associated with mutations in *LRP1B, TP53*, and *KEAP1* (**Supplementary Figure S2E**). Then, we used the “Subgroup Analysis” module to calculate the P interaction between the *STK11* mutation and other gene mutations that were correlated with *STK11*. It turned out that the *STK11* mutation significantly interacted with *KEAP1* on the Samstein_2018 dataset (P interaction = 0.04, **Figure 2E**).

#### 3. Potential exploration in the biological mechanism

The correlation of the *STK11* mutation with immunogenicity and immune signatures was studied with the help of “Immunogenicity Analysis” and “Microenvironment Analysis” modules. In the TCGA-LUAD dataset, patients with the *STK11* mutation had significantly lower levels of Tertiary Lymphoid Structure (TLS) score, Cytolytic Activity (CYT) score, ssGSEA score of IL-6 pathway, antigen processing MHC-II, TGF-beta pathway, PD-1 pathway, and interferon gamma pathway (P < 0.05, **Figure 2F-G**), suggesting that worse prognosis of *STK11* mutation in LUAD could be due to worse immune activation and reaction, which might provide some clues for further mechanism study.

### Application 2: To identify a biomarker for ICIs: predictive and/or prognostic

When a biomarker is associated with survival in patients treated with ICIs, the interaction effect analysis between biomarker and treatment group is critical to determine whether it is a predictive biomarker ^12^. In the application 1, we learnt that *STK11* was an independent biomarker in ICIs cohorts. Here, we further explored whether *STK11* was a predictive or prognostic biomarker using the Gandara_2018 dataset which was a good example to illustrate the necessity of interaction effect analysis for identifying predictive and/or prognostic biomarkers. Gandara_2018 dataset ^14^ consists of two randomized control studies (POPLAR and OAK) in NSCLC (atezolizumab as the intervention arm and docetaxel as the control arm). In the intervention arm, patients with the *STK11* mutation had shorter OS than those with wild type (Log-rank P = 0.005, **Figure 3A**); whereas there were similar results in the control arm (Log-rank P = 0.001, **Figure 3A**). Furthermore, P for the interaction of 0.96 suggests that the difference between mutation and wild-type groups of *STK11* was consistent in immunotherapy and docetaxel arms. Therefore, the *STK11* mutation was a prognostic biomarker but not a predictive biomarker especially for ICIs in LUAD.

**Figure 3.**
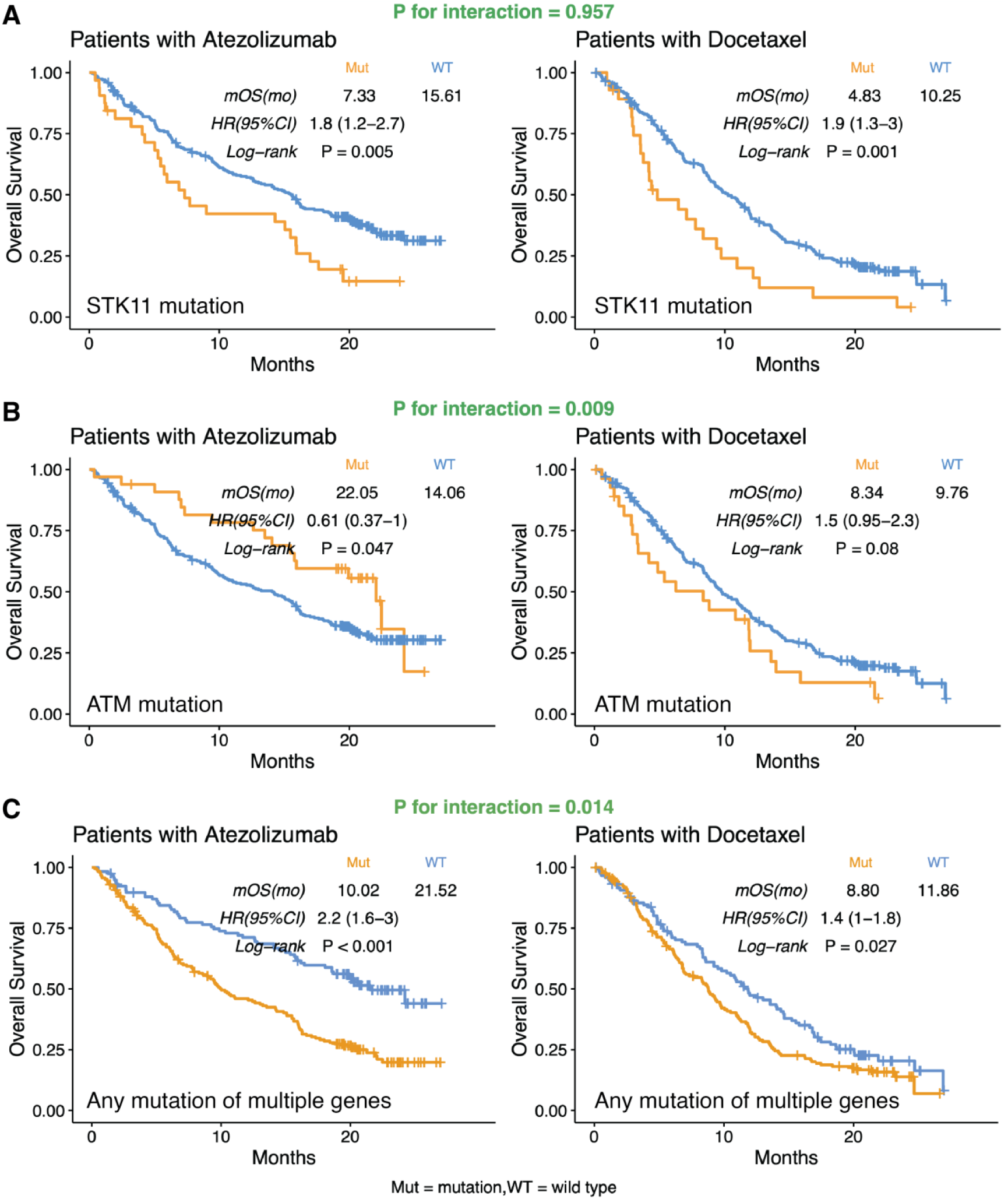
The interaction analysis between gene mutations and treatment on the Gandara_2018 dataset. **(A)** The Kaplan-Meier curve of *STK11* mutation for OS in patients with Atezolizumab and Docetaxel, respectively. **(B)** The Kaplan-Meier curve of *ATM* mutation for overall survival in patients with Atezolizumab and Docetaxel, respectively. **(C)** The Kaplan-Meier curve of any mutation of *STK11, TP53, KEAP1, EGFR, ERBB4*, and *NTRK3* for OS in patients with Atezolizumab and Docetaxel, respectively.

We also provided another two examples to demonstrate the identification of predictive and/or prognostic biomarkers for ICIs: (1) *ATM* mutation. *ATM* mutation was positively significantly associated with OS only in the intervention arm (Log-rank P = 0.05, HR = 0.61, 95% CI = 0.37 - 1.00), not in the control arm (Log-rank P = 0.08, HR = 1.50, 95% CI = 0.95 - 2.30; P for interaction = 0.009, **Figure 3B**), suggesting *ATM* mutation is more likely to be a predictive biomarker for ICIs in these patients; and (2) a multi-gene biomarker, including *STK11, TP53, KEAP1, EGFR, ERBB4*, and *NTRK3*. The “biomarker-WT” group was defined as patients without non-synonymous mutations of any of those genes, while the “biomarker-MUT” group was defined as patients with non-synonymous mutation in any of those genes. In both intervention and control arms, biomarker-WT patients had better prognoses than biomarker-MUT patients (**Figure 3C**). However, the benefit of prognosis between biomarker-WT and biomarker-MUT was greater in the intervention arm than in the control arm (HR: 2.2 vs 1.4, P for interaction = 0.01, **Figure 3C**). Apparently, this multi-gene biomarker could function as both predictive and prognostic for LUAD patients. Therefore, the “Interaction Analysis” can clearly demonstrate the predictive and prognostic nature of a biomarker.

## Discussion

Discovering and identifying biomarkers for ICIs is an urgent need and one of the prerequisites for effective immunotherapy, and demonstrating whether a biomarker is predictive and/or prognostic could be also important for treatment choices. Therefore, we developed IMPACT, the most systematic analysis web-based platform for exploring ICIs prediction biomarkers, to validate previous findings and advance the discovery of novel biomarkers.

As a web server, IMPACT attempts to provide multiple unique and helpful features. It curates a more comprehensive resource of ICIs data than other tools, allows users to upload and analyze their own data, and timely updates new available data, to provide a more robust and applicable tool for customized scenarios. To thoroughly investigate biomarkers for ICIs, IMPACT integrates and optimizes conventional functions of biomarker exploration tools to simplify and visualize bioinformatic analyses, including univariable and multivariable analyses, meta-analysis, subgroup analysis, and association analysis between biomarkers and immune signatures.

Intriguingly, IMPACT exclusively provides the interaction effect analysis to unveil the interaction between biomarkers and genes of interest to discover co-mutation biomarkers with better performance, which can be more efficient. In a previous study, the predictive effects of *KRAS* depended on the presence of other co-mutation genes. *KRAS*/*TP53* co-mutations favored survival with ICIs but *KRAS*/*STK11* co-mutation had the opposite results ^15^. In addition, it was also reported that the positive effects of *TP53* mutation might be neutralized by *EGFR* mutation so that *TP53*/*EGFR* co-mutation had a worse prognosis for ICIs treatment than *TP53* mutation, therefore, ICIs may not be the first choice for *TP53*/*EGFR* patients ^16^. The above results could be easily obtained on IMPACT. Different from the other tool that users need to manually select two genes of interest to analyze the effect of co-mutation, IMPACT allows the users to generate a list of target genes (up to 50) and automatically analyze all the pair-wise co-mutations, to provide comprehensive profile of co-mutation for the further analyses. Moreover, growing evidence suggests that different mutation variants of the same gene could be associated with different outcomes, which might be a reflection of different biological mechanisms. Therefore, the function of selecting a certain type of mutation in IMPACT can advance finding clinically meaningful biomarkers. As reported in kidney cancer, patients with *PBRM1*-truncating mutation, a loss of function mutation, had significantly longer OS than those with *PBRM1* non-truncating mutation ^17^. The same result was obtained by IMPACT (**Supplementary Figure S1A-B**), while none of the existing tools could customize the mutation variant types of interest in such details.

Moreover, the interaction effect analysis also unveils the interaction between biomarkers and treatment groups to determine the nature of a biomarker, to be prognostic and/or predictive, therefore efficiently screening the beneficiaries for ICIs treatments. *ATM* mutation has been reported as a predictive biomarker by analyzing the ICIs databases of LUAD ^18^. However, it was undefined whether it is a specific predictive biomarker for ICIs. With the interaction analysis of *ATM* mutation and the ICIs treatment using IMPACT, *ATM* mutation was identified as a predictive biomarker for ICIs as it was not associated with OS in the non-ICIs arm of LUAD patients and the P for interaction is significant. Interestingly, a previous study had identified the *STK11* alterations as a major driver of poor prognosis for ICIs in the *KRAS*-mutant LUAD ^6^. However, according to the interaction analysis using IMPACT, *STK11* mutation was a prognosis biomarker for both ICIs and chemotherapy, but not a predictive biomarker specified for ICIs, which was supported by recent reports ^19^. Therefore, the conclusion might be drawn that the interaction analyses in IMPACT are essential for exploring biomarkers which fills the functional blank of existing tools.

Furthermore, the interaction between biomarkers and other clinical characteristics can distinguish the treatment outcomes with more precision stratification, which can also be performed by IMPACT. For continuous variables, such as gene expression or TMB, the cut-off value selection can significantly affect the analysis results. In a blood-based TMB (bTMB) study by Gandara *et al*. ^14^, a higher bTMB derived by a predefined cut-off was not associated with OS, but the forest plot of HR based on various bTMB cut-offs showed that bTMB tended to be negatively associated with OS, which inspired another valuable work of using maximum somatic allele frequency (MSAF) modified bTMB to predict ICIs outcome ^20^. However, the existing tools only allow one predefined cut-off in one analysis, impeding the observation of the variation trend of association across different cut-offs. Hopefully, the function of studying continuous variables across different cut-offs provided by IMPACT might also contribute to finding new potential biomarkers.

In summary, IMPACT is a user-friendly platform conveying a comprehensive resource for exploring immunotherapeutic predictive and cancer prognostic biomarkers, which eases bioinformatic analyses for researchers. Using IMPACT, users can deeply explore predictive and/or prognostic biomarkers, relevant interaction effects, and potential biological mechanisms. With long-term support, continuous upgrade and optimization, IMPACT has the potential to become a popular tool to facilitate immunotherapy research.

## Acknowledgments

We thank all the volunteer patients, staffs and researchers in TCGA, GEO and relevant published literature for their contribution to the research data.

## Author Contribution

Conceptualization: YL, WX, JZ, SC, ZW; Data curation: YL, YZ, WX, JZ, YD, CX, ML, KL, HL, JD; Software: WX, YW; Writing-original draft: YL, YZ, WX, JZ, YW, GW, XZ; Writing-review and editing: all authors; Supervision: SC, JW, ZW; Funding acquisition: JW, ZW

