## Supplemental Figures for "IMPACT: a web server for exploring immunotherapeutic predictive and cancer prognostic biomarkers"

**Table S1. The description of ICIs datasets**

| Data source | Study ID | Treatment | Cancer type | PMID | Sample size |
|---|---|---|---|---|---|
| Public | Allen_2015 | anti-CTLA-4 | Melanoma | 26359337 | 110 |
| Public | Hugo_2016 | anti-PD-1/L1 | Melanoma | 26997480 | 38 |
| Public | Synder_2014 | anti-CTLA-4 | Melanoma | 25409260 | 64 |
| Public | Liu_2019 | anti-PD-1/L1 | Melanoma | 31792460 | 144 |
| Public | Riaz_2017 | anti-PD-1/L1,anti-CTLA-4 | Melanoma | 29033130 | 73 |
| Public | Gide_2019 | anti-PD-1/L1,anti-CTLA-4 | Melanoma | 30753825 | 105 |
| Public | Jerby_Arnon_2018 | anti-PD-1/L1 | Melanoma | 30388455 | 104 |
| Public | Hellmann_2018 | anti-PD-1/L1,anti-CTLA-4 | LUAD | 29657128 | 59 |
| Public | Hellmann_2018 | anti-PD-1/L1,anti-CTLA-4 | LUSC | 29657128 | 16 |
| Public | Gandara_2018 | anti-PD-1/L1 | LUAD | 30082870 | 598 |
| Public | Gandara_2018 | anti-PD-1/L1 | LUSC | 30082870 | 255 |
| Public | Rizvi_2015 | anti-PD-1/L1 | LUAD | 25765070 | 29 |
| Public | Rizvi_2018 | anti-PD-1/L1,anti-CTLA-4 | LUAD | 29337640 | 186 |
| Public | Rizvi_2018 | anti-PD-1/L1,anti-CTLA-4 | LUSC | 29337640 | 34 |
| Public | GSE135222 | anti-PD-1/L1 | NSCLC | 31537801 | 60 |
| Public | GSE136961 | anti-PD-1/L1 | NSCLC | 31959763 | 21 |
| Public | Samstein_2018 | anti-PD-1/L1,anti-CTLA-4 | BLCA | 30643254 | 215 |
| Public | Samstein_2018 | anti-PD-1/L1,anti-CTLA-4 | BRCA | 30643254 | 44 |
| Public | Samstein_2018 | anti-PD-1/L1,anti-CTLA-4 | COAD | 30643254 | 110 |
| Public | Samstein_2018 | anti-PD-1/L1,anti-CTLA-4 | ESCA | 30643254 | 126 |
| Public | Samstein_2018 | anti-PD-1/L1,anti-CTLA-4 | Glioma | 30643254 | 117 |
| Public | Samstein_2018 | anti-PD-1/L1,anti-CTLA-4 | HNSC | 30643254 | 139 |
| Public | Samstein_2018 | anti-PD-1/L1,anti-CTLA-4 | KIRC | 30643254 | 151 |

| | | | | | |
|---|---|---|---|---|---|
| Public | Samstein_2018 | anti-PD-1/L1,anti-CTLA-4 | Melanoma | 30643254 | 320 |
| Public | Samstein_2018 | anti-PD-1/L1,anti-CTLA-4 | LUAD | 30643254 | 271 |
| Public | Samstein_2018 | anti-PD-1/L1,anti-CTLA-4 | LUSC | 30643254 | 45 |
| Public | Miao_2018 | anti-PD-1/L1,anti-CTLA-4 | BLCA | 30150660 | 27 |
| Public | Miao_2018 | anti-PD-1/L1,anti-CTLA-4 | Melanoma | 30150660 | 151 |
| Public | Miao_2018 | anti-PD-1/L1,anti-CTLA-4 | LUAD | 30150660 | 50 |
| Public | Motzer_2020 | anti-PD-1/L1 | KIRC | 32895571 | 886 |
| Public | Checkmate009 | anti-PD-1/L1 | KIRC | 32472114 | 35 |
| Public | Checkmate010 | anti-PD-1/L1 | KIRC | 32472114 | 168 |
| Public | Checkmate025 | anti-PD-1/L1 | KIRC | 32472114 | 803 |
| Public | Mariathasan_2018 | anti-PD-1/L1 | BLCA | 29443960 | 348 |
| Public | GSE176307 | anti-PD-1/L1 | BLCA | 34294892 | 89 |
| In-house | Bai_2020 | anti-PD-1/L1 | LUAD | 33303576 | 54 |
| In-house | Bai_2020 | anti-PD-1/L1 | LUSC | 33303576 | 25 |
| In-house | NCC-ICIs | anti-PD-1/L1 | LUAD | / | 30 |
| In-house | NCC-ICIs | anti-PD-1/L1 | LUSC | / | 19 |
| In-house | Wang_2019 | anti-PD-1/L1 | LUAD | 30816954 | 31 |
| In-house | Wang_2019 | anti-PD-1/L1 | LUSC | 30816954 | 19 |
| Public | Subudhi_2020 | anti-CTLA-4 | CRPC | 32238575 | 30 |
| Public | Margolis_2018 | anti-PD-1/L1 | KIRC | 29301960 | 35 |
| Public | Zhao_2019 | anti-PD-1/L1 | Glioma | 30742119 | 42 |

**Table S2. The association of driver mutation or co-mutation with PFS or OS in each ICIs dataset.**

| Cohort | Cancer | Gene | Desciption | NoOfPts[1] | NoOfMut[2] | PFS_m_mut[3] | PFS_m_wt[4] | PFS_HR95CI[5] | PFS_Cox_P[6] | OS_m_mut[7] | OS_m_wt[8] | OS_HR95CI[9] | OS_Cox_P[10] |
|---|---|---|---|---|---|---|---|---|---|---|---|---|---|
| Hellmann_2018 (WES) | LUAD | TP53 | MUT/WT | 59 | 29 | 7.98 | 7.56 | 0.6 (0.32-1.13) | 0.11 | | | | |
| Hellmann_2018 (WES) | LUAD | KRAS | MUT/WT | 59 | 23 | 7.24 | 7.82 | 0.93 (0.49-1.76) | 0.82 | | | | |
| Hellmann_2018 (WES) | LUAD | STK11 | MUT/WT | 59 | 12 | 6.51 | 7.82 | 1.28 (0.58-2.79) | 0.54 | | | | |
| Hellmann_2018 (WES) | LUAD | KEAP1 | MUT/WT | 59 | 11 | 7.95 | 7.82 | 0.65 (0.27-1.54) | 0.33 | | | | |
| Hellmann_2018 (WES) | LUAD | EGFR | MUT/WT | 59 | 10 | 7.79 | 7.82 | 1.2 (0.55-2.6) | 0.65 | | | | |
| Hellmann_2018 (WES) | LUAD | TP53#KRAS | MUT/WT | 59 | 11 | NA | 7.75 | 0.35 (0.12-0.99) | 0.048 | | | | |
| Hellmann_2018 (WES) | LUAD | TP53#STK11 | MUT/WT | 59 | 4 | 2.1 | 7.82 | 0.82 (0.2-3.44) | 0.79 | | | | |
| Hellmann_2018 (WES) | LUAD | TP53#KEAP1 | MUT/WT | 59 | 5 | NA | 7.75 | 0.17 (0.02-1.25) | 0.082 | | | | |
| Hellmann_2018 (WES) | LUAD | TP53#EGFR | MUT/WT | 59 | 4 | 4.81 | 7.82 | 1.37 (0.42-4.47) | 0.6 | | | | |
| Hellmann_2018 (WES) | LUAD | KRAS#STK11 | MUT/WT | 59 | 8 | 2.3 | 7.95 | 1.6 (0.62-4.11) | 0.33 | | | | |
| Hellmann_2018 (WES) | LUAD | KRAS#KEAP1 | MUT/WT | 59 | 5 | 6.51 | 7.82 | 0.86 (0.26-2.79) | 0.8 | | | | |
| Hellmann_2018 | LUA | KRAS#EGFR | MUT/WT | 59 | 2 | 7.98 | 7.75 | 0.42 | 0.4 | | | | |

| | | | | | | | | | | | | | |
|---|---|---|---|---|---|---|---|---|---|---|---|---|---|
| (WES) | D | | | | | | | (0.06-3.1) | | | | | |
| Hellmann_2018 (WES) | LUAD | STK11#KEAP1 | MUT/WT | 59 | 6 | 4.5 | 7.82 | 1.74 (0.68-4.5) | 0.25 | | | | |
| Hellmann_2018 (WES) | LUAD | STK11#EGFR | All WT | 59 | 0 | | | | | | | | |
| Hellmann_2018 (WES) | LUAD | KEAP1#EGFR | MUT/WT | 59 | 2 | NA | 7.75 | 0 (0-Inf) | 1 | | | | |
| Hellmann_2018 (WES) | LUAD | TP53#KRAS#STK11#KEAP1#EGFR | All WT | 59 | 0 | | | | | | | | |
| Gandara_2018 (FM) | LUAD | TP53 | MUT/WT | 301 | 133 | 2.69 | 2.79 | 1.08 (0.85-1.38) | 0.52 | 11.01 | 15.9 | 1.39 (1.04-1.84) | 0.024 |
| Gandara_2018 (FM) | LUAD | KRAS | MUT/WT | 301 | 33 | 1.51 | 2.78 | 1.27 (0.87-1.84) | 0.22 | 10.61 | 15.34 | 1.42 (0.92-2.18) | 0.11 |
| Gandara_2018 (FM) | LUAD | STK11 | MUT/WT | 301 | 32 | 1.41 | 2.79 | 1.54 (1.04-2.26) | 0.029 | 7.33 | 15.61 | 1.81 (1.19-2.74) | 0.0052 |
| Gandara_2018 (FM) | LUAD | KEAP1 | MUT/WT | 301 | 45 | 1.41 | 2.79 | 1.58 (1.14-2.2) | 0.006 | 7.06 | 15.97 | 2 (1.4-2.86) | 0.00016 |
| Gandara_2018 (FM) | LUAD | EGFR | MUT/WT | 301 | 38 | 2.53 | 2.79 | 1.31 (0.92-1.87) | 0.14 | 10.45 | 15.47 | 1.29 (0.85-1.95) | 0.23 |
| Gandara_2018 (FM) | LUAD | TP53#KRAS | MUT/WT | 301 | 16 | 5.17 | 2.73 | 0.8 (0.47-1.37) | 0.42 | 16.03 | 14.49 | 0.95 (0.5-1.8) | 0.87 |
| Gandara_2018 (FM) | LUAD | TP53#STK11 | MUT/WT | 301 | 14 | 1.51 | 2.73 | 0.89 (0.49-1.63) | 0.71 | 6.85 | 15.05 | 1.55 (0.84-2.85) | 0.16 |
| Gandara_2018 (FM) | LUAD | TP53#KEAP1 | MUT/WT | 301 | 24 | 1.49 | 2.73 | 1.31 (0.85-2) | 0.22 | 10.61 | 15.47 | 1.43 (0.88-2.32) | 0.15 |

| | | | | | | | | | | | | | |
|---|---|---|---|---|---|---|---|---|---|---|---|---|---|
| Gandara_2018 (FM) | LUAD | TP53#EGFR | MUT/WT | 301 | 25 | 2.69 | 2.74 | 1.22 (0.8-1.88) | 0.35 | 9.46 | 15.34 | 1.3 (0.79-2.14) | 0.3 |
| Gandara_2018 (FM) | LUAD | KRAS#STK11 | MUT/WT | 301 | 4 | 1.17 | 2.76 | 13.08 (4.56-37.46) | 1.70E-06 | 4.25 | 15.05 | 3.39 (1.26-9.17) | 0.016 |
| Gandara_2018 (FM) | LUAD | KRAS#KEAP1 | MUT/WT | 301 | 8 | 1.41 | 2.76 | 1.29 (0.61-2.74) | 0.51 | 8.84 | 15.34 | 2.07 (0.97-4.41) | 0.06 |
| Gandara_2018 (FM) | LUAD | KRAS#EGFR | MUT/WT | 301 | 1 | 1.31 | 2.73 | 5.25 (0.73-37.93) | 0.1 | 7.82 | 15.05 | 2.35 (0.33-16.85) | 0.4 |
| Gandara_2018 (FM) | LUAD | STK11#KEAP1 | MUT/WT | 301 | 11 | 1.28 | 2.79 | 2.9 (1.58-5.33) | 6.00E-04 | 5.29 | 15.47 | 2.69 (1.46-4.96) | 0.0016 |
| Gandara_2018 (FM) | LUAD | STK11#EGFR | MUT/WT | 301 | 2 | 10.35 | 2.73 | 0.28 (0.04-1.97) | 0.2 | 15.93 | 14.82 | 0.58 (0.08-4.12) | 0.58 |
| Gandara_2018 (FM) | LUAD | KEAP1#EGFR | MUT/WT | 301 | 4 | 7.21 | 2.73 | 0.6 (0.19-1.88) | 0.38 | 14.18 | 14.98 | 1.15 (0.37-3.61) | 0.81 |
| Gandara_2018 (FM) | LUAD | TP53#KRAS#STK11#KEAP1#EGFR | All WT | 301 | 0 | | | | | | | | |
| Rizvi_2015 (WES) | LUAD | TP53 | MUT/WT | 29 | 12 | 14.7 | 10.4 | 0.25 (0.05-1.33) | 0.1 | | | | |
| Rizvi_2015 (WES) | LUAD | KRAS | MUT/WT | 29 | 8 | 14.6 | 10.4 | 0.24 (0.04-1.41) | 0.11 | | | | |
| Rizvi_2015 (WES) | LUAD | STK11 | MUT/WT | 29 | 5 | 9.5 | 12.6 | 3.14 (0.68-14.41) | 0.14 | | | | |
| Rizvi_2015 (WES) | LUAD | KEAP1 | MUT/WT | 29 | 4 | 14.6 | 10.4 | 1.09 (0.24-4.91) | 0.91 | | | | |
| Rizvi_2015 (WES) | LUA | EGFR | MUT/WT | 29 | 2 | NA | 12.6 | NA (NA-NA) | NA | | | | |

| | | | | | | | | | |
|---|---|---|---|---|---|---|---|---|---|
| | D | | | | | | | | |
| Rizvi_2015 (WES) | LUAD | TP53#KRAS | MUT/WT | 29 | 4 | 14.7 | 10.4 | 0.18 (0.02-1.54) | 0.12 |
| Rizvi_2015 (WES) | LUAD | TP53#STK11 | MUT/WT | 29 | 1 | NA | 12.6 | NA (NA-NA) | NA |
| Rizvi_2015 (WES) | LUAD | TP53#KEAP1 | MUT/WT | 29 | 2 | 14.7 | 10.4 | 0.5 (0.06-4.36) | 0.53 |
| Rizvi_2015 (WES) | LUAD | TP53#EGFR | MUT/WT | 29 | 2 | NA | 12.6 | NA (NA-NA) | NA |
| Rizvi_2015 (WES) | LUAD | KRAS#STK11 | MUT/WT | 29 | 1 | 14.6 | 11.5 | 0.9 (0.1-7.93) | 0.92 |
| Rizvi_2015 (WES) | LUAD | KRAS#KEAP1 | MUT/WT | 29 | 2 | 14.65 | 10.4 | 0.62 (0.11-3.43) | 0.59 |
| Rizvi_2015 (WES) | LUAD | KRAS#EGFR | All WT | 29 | 0 | | | | |
| Rizvi_2015 (WES) | LUAD | STK11#KEAP1 | MUT/WT | 29 | 2 | 9.4 | 12.6 | 2.13 (0.39-11.57) | 0.38 |
| Rizvi_2015 (WES) | LUAD | STK11#EGFR | All WT | 29 | 0 | | | | |
| Rizvi_2015 (WES) | LUAD | KEAP1#EGFR | All WT | 29 | 0 | | | | |
| Rizvi_2015 (WES) | LUAD | TP53#KRAS#STK11#KEAP1#EGFR | All WT | 29 | 0 | | | | |
| Rizvi_2018 (MSK-IMPACT) | LUAD | TP53 | MUT/WT | 185 | 102 | 4.2 | 2.57 | 0.67 (0.49-0.92) | 0.014 |

| | | | | | | | | | |
|---|---|---|---|---|---|---|---|---|---|
| Rizvi_2018 (MSK-IMPACT) | LUAD | KRAS | MUT/WT | 185 | 77 | 3.77 | 3.3 | 0.9 (0.65-1.23) | 0.5 |
| Rizvi_2018 (MSK-IMPACT) | LUAD | STK11 | MUT/WT | 185 | 49 | 2.5 | 4 | 1.26 (0.89-1.78) | 0.2 |
| Rizvi_2018 (MSK-IMPACT) | LUAD | KEAP1 | MUT/WT | 185 | 41 | 2.8 | 3.5 | 0.98 (0.67-1.42) | 0.9 |
| Rizvi_2018 (MSK-IMPACT) | LUAD | EGFR | MUT/WT | 185 | 25 | 3.07 | 3.6 | 1.53 (0.97-2.41) | 0.068 |
| Rizvi_2018 (MSK-IMPACT) | LUAD | TP53#KRAS | MUT/WT | 185 | 27 | 5.47 | 3.07 | 0.49 (0.29-0.81) | 0.0059 |
| Rizvi_2018 (MSK-IMPACT) | LUAD | TP53#STK11 | MUT/WT | 185 | 17 | 3.3 | 3.5 | 0.98 (0.57-1.7) | 0.95 |
| Rizvi_2018 (MSK-IMPACT) | LUAD | TP53#KEAP1 | MUT/WT | 185 | 24 | 3.38 | 3.43 | 0.91 (0.57-1.48) | 0.71 |
| Rizvi_2018 (MSK-IMPACT) | LUAD | TP53#EGFR | MUT/WT | 185 | 18 | 3.4 | 3.43 | 1.35 (0.8-2.26) | 0.26 |
| Rizvi_2018 (MSK-IMPACT) | LUAD | KRAS#STK11 | MUT/WT | 185 | 28 | 2.24 | 3.6 | 1.24 (0.8-1.9) | 0.33 |
| Rizvi_2018 (MSK-IMPACT) | LUAD | KRAS#KEAP1 | MUT/WT | 185 | 17 | 2.27 | 3.5 | 0.96 (0.55-1.67) | 0.89 |
| Rizvi_2018 (MSK-IMPACT) | LUAD | KRAS#EGFR | MUT/WT | 185 | 2 | 2.05 | 3.5 | 2.84 (0.7-11.58) | 0.15 |
| Rizvi_2018 (MSK-IMPACT) | LUAD | STK11#KEAP1 | MUT/WT | 185 | 24 | 2.18 | 3.6 | 1.1 (0.69-1.75) | 0.68 |
| Rizvi_2018 (MSK-IMPACT) | LUAD | STK11#EGFR | MUT/WT | 185 | 2 | 2.88 | 3.5 | 1.7 (0.42-6.93) | 0.46 |

| | | | | | | | | | | | | | |
|---|---|---|---|---|---|---|---|---|---|---|---|---|---|
| Rizvi_2018 (MSK-IMPACT) | LUAD | KEAP1#EGFR | MUT/WT | 185 | 1 | 2.47 | 3.5 | 2 (0.28-14.43) | 0.49 | | | | |
| Rizvi_2018 (MSK-IMPACT) | LUAD | TP53#KRAS#STK11#KEAP1#EGFR | All WT | 185 | 0 | | | | | | | | |
| Samstein_2018 (IMPACT) | LUAD | TP53 | MUT/WT | 266 | 157 | | | | | 11 | 14 | 1.17 (0.85-1.6) | 0.33 |
| Samstein_2018 (IMPACT) | LUAD | KRAS | MUT/WT | 266 | 109 | | | | | 12 | 12 | 0.96 (0.7-1.31) | 0.79 |
| Samstein_2018 (IMPACT) | LUAD | STK11 | MUT/WT | 266 | 66 | | | | | 6 | 13 | 1.47 (1.05-2.06) | 0.027 |
| Samstein_2018 (IMPACT) | LUAD | KEAP1 | MUT/WT | 266 | 60 | | | | | 9 | 13 | 1.46 (1.02-2.09) | 0.039 |
| Samstein_2018 (IMPACT) | LUAD | EGFR | MUT/WT | 266 | 41 | | | | | 13 | 12 | 0.95 (0.61-1.47) | 0.81 |
| Samstein_2018 (IMPACT) | LUAD | TP53#KRAS | MUT/WT | 266 | 46 | | | | | 12 | 12 | 0.95 (0.63-1.45) | 0.82 |
| Samstein_2018 (IMPACT) | LUAD | TP53#STK11 | MUT/WT | 266 | 26 | | | | | 4.5 | 13 | 1.47 (0.9-2.39) | 0.13 |
| Samstein_2018 (IMPACT) | LUAD | TP53#KEAP1 | MUT/WT | 266 | 33 | | | | | 12 | 12 | 1.06 (0.66-1.72) | 0.8 |
| Samstein_2018 (IMPACT) | LUAD | TP53#EGFR | MUT/WT | 266 | 30 | | | | | 13 | 12 | 1.01 (0.62-1.65) | 0.96 |
| Samstein_2018 (IMPACT) | LUAD | KRAS#STK11 | MUT/WT | 266 | 38 | | | | | 8 | 12 | 1.31 (0.87-1.99) | 0.2 |
| Samstein_2018 | LUA | KRAS#KEAP1 | MUT/WT | 266 | 30 | | | | | 4 | 13 | 1.96 | 0.003 |

| | | | | | | | | | | | | | |
|---|---|---|---|---|---|---|---|---|---|---|---|---|---|
| (IMPACT) | D | | | | | | | | | | | (1.26-3.05) | |
| Samstein_2018 (IMPACT) | LUAD | KRAS#EGFR | MUT/WT | 266 | 1 | | | | | 13 | 12 | 1.33 (0.19-9.51) | 0.78 |
| Samstein_2018 (IMPACT) | LUAD | STK11#KEAP1 | MUT/WT | 266 | 36 | | | | | 4 | 13 | 2.09 (1.39-3.15) | 0.00042 |
| Samstein_2018 (IMPACT) | LUAD | STK11#EGFR | MUT/WT | 266 | 1 | | | | | NA | 12 | 0 (0-Inf) | 0.99 |
| Samstein_2018 (IMPACT) | LUAD | KEAP1#EGFR | MUT/WT | 266 | 1 | | | | | 10 | 12 | 1.67 (0.23-11.97) | 0.61 |
| Samstein_2018 (IMPACT) | LUAD | TP53#KRAS#STK11#KEAP1#EGFR | All WT | 266 | 0 | | | | | | | | |
| Miao_2018 (WES) | LUAD | TP53 | MUT/WT | 50 | 28 | 5.22 | 5.13 | 1.05 (0.55-2.01) | 0.89 | 13.77 | 27.27 | 1.19 (0.46-3.1) | 0.72 |
| Miao_2018 (WES) | LUAD | KRAS | MUT/WT | 50 | 16 | 7.65 | 3.75 | 0.52 (0.25-1.09) | 0.083 | 27.27 | 12.8 | 0.61 (0.23-1.65) | 0.33 |
| Miao_2018 (WES) | LUAD | STK11 | MUT/WT | 50 | 6 | 1.8 | 5.87 | 2.4 (0.92-6.27) | 0.073 | 4.4 | 15.73 | 3.48 (1.12-10.83) | 0.031 |
| Miao_2018 (WES) | LUAD | KEAP1 | MUT/WT | 50 | 11 | 5.67 | 4.2 | 0.8 (0.35-1.83) | 0.6 | 12.1 | 27.27 | 1.9 (0.69-5.25) | 0.22 |
| Miao_2018 (WES) | LUAD | EGFR | MUT/WT | 50 | 14 | 1.92 | 6.57 | 2.76 (1.38-5.52) | 0.004 | 11.3 | 27.27 | 1.63 (0.6-4.42) | 0.34 |
| Miao_2018 (WES) | LUAD | TP53#KRAS | MUT/WT | 50 | 6 | 14.7 | 4.18 | 0.39 (0.12-1.27) | 0.12 | NA | 15.73 | 0.45 (0.1-2) | 0.3 |
| Miao_2018 (WES) | LUAD | TP53#STK11 | MUT/WT | 50 | 2 | 4.72 | 4.77 | 1.7 (0.41-7.14) | 0.47 | 8.1 | 15.73 | 0.98 (0.13-7.44) | 0.99 |

| | | | | | | | | | | | | | |
|---|---|---|---|---|---|---|---|---|---|---|---|---|---|
| Miao_2018 (WES) | LUAD | TP53#KEAP1 | MUT/WT | 50 | 5 | 5.67 | 4.2 | 0.9 (0.32-2.56) | 0.85 | 13.77 | 27.27 | 1.45 (0.41-5.09) | 0.56 |
| Miao_2018 (WES) | LUAD | TP53#EGFR | MUT/WT | 50 | 10 | 1.88 | 6.37 | 2.67 (1.24-5.75) | 0.012 | 11.3 | 27.27 | 1.89 (0.66-5.4) | 0.23 |
| Miao_2018 (WES) | LUAD | KRAS#STK11 | MUT/WT | 50 | 2 | 1.4 | 5.67 | 13.71 (2.74-68.48) | 0.0014 | 2.52 | 15.73 | 27.37 (4.45-168.33) | 0.00036 |
| Miao_2018 (WES) | LUAD | KRAS#KEAP1 | MUT/WT | 50 | 6 | 4.23 | 5.67 | 0.93 (0.33-2.64) | 0.89 | 12.93 | 27.27 | 1.82 (0.58-5.7) | 0.31 |
| Miao_2018 (WES) | LUAD | KRAS#EGFR | All WT | 50 | 0 | | | | | | | | |
| Miao_2018 (WES) | LUAD | STK11#KEAP1 | MUT/WT | 50 | 4 | 1.8 | 5.87 | 3.02 (0.89-10.23) | 0.076 | 4.18 | 27.27 | 15.08 (2.96-76.71) | 0.0011 |
| Miao_2018 (WES) | LUAD | STK11#EGFR | MUT/WT | 50 | 1 | 1.33 | 5.67 | 9.29 (1.09-79.53) | 0.042 | NA | 15.73 | 0 (0-Inf) | 1 |
| Miao_2018 (WES) | LUAD | KEAP1#EGFR | MUT/WT | 50 | 1 | NA | 4.77 | 0 (0-Inf) | 1 | NA | 15.73 | 0 (0-Inf) | 1 |
| Miao_2018 (WES) | LUAD | TP53#KRAS#STK11#KEAP1#EGFR | All WT | 50 | 0 | | | | | | | | |
| Bai_2020 (WES) | LUAD | TP53 | MUT/WT | 50 | 25 | 2.13 | 2.03 | 0.64 (0.36-1.16) | 0.14 | | | | |
| Bai_2020 (WES) | LUAD | KRAS | MUT/WT | 50 | 6 | 5.82 | 2.1 | 0.72 (0.31-1.71) | 0.46 | | | | |
| Bai_2020 (WES) | LUAD | STK11 | MUT/WT | 50 | 1 | 4.23 | 2.1 | 0.84 (0.11-6.17) | 0.87 | | | | |
| Bai_2020 (WES) | LUA | KEAP1 | MUT/WT | 50 | 2 | 7.6 | 2.1 | 0.58 | 0.45 | | | | |

| | | | | | | | | | |
|---|---|---|---|---|---|---|---|---|---|
| | D | | | | | | | (0.14-2.4) | |
| Bai_2020 (WES) | LUAD | EGFR | MUT/WT | 50 | 9 | 2.1 | 2.1 | 1.53 (0.72-3.25) | 0.27 |
| Bai_2020 (WES) | LUAD | TP53#KRAS | MUT/WT | 50 | 2 | 5.95 | 2.1 | 0.91 (0.22-3.8) | 0.9 |
| Bai_2020 (WES) | LUAD | TP53#STK11 | All WT | 50 | 0 | | | | |
| Bai_2020 (WES) | LUAD | TP53#KEAP1 | MUT/WT | 50 | 1 | 11.1 | 2.1 | 0.43 (0.06-3.18) | 0.41 |
| Bai_2020 (WES) | LUAD | TP53#EGFR | MUT/WT | 50 | 4 | 1.93 | 2.1 | 1.55 (0.55-4.37) | 0.41 |
| Bai_2020 (WES) | LUAD | KRAS#STK11 | All WT | 50 | 0 | | | | |
| Bai_2020 (WES) | LUAD | KRAS#KEAP1 | MUT/WT | 50 | 1 | 4.1 | 2.1 | 0.89 (0.12-6.53) | 0.91 |
| Bai_2020 (WES) | LUAD | KRAS#EGFR | MUT/WT | 50 | 1 | 4.1 | 2.1 | 0.89 (0.12-6.53) | 0.91 |
| Bai_2020 (WES) | LUAD | STK11#KEAP1 | All WT | 50 | 0 | | | | |
| Bai_2020 (WES) | LUAD | STK11#EGFR | MUT/WT | 50 | 1 | 4.23 | 2.1 | 0.84 (0.11-6.17) | 0.87 |
| Bai_2020 (WES) | LUAD | KEAP1#EGFR | MUT/WT | 50 | 1 | 4.1 | 2.1 | 0.89 (0.12-6.53) | 0.91 |
| Bai_2020 (WES) | LUAD | TP53#KRAS#STK11#KEAP1#EGFR | All WT | 50 | 0 | | | | |

| | | | | | | | | | | | | | |
|---|---|---|---|---|---|---|---|---|---|---|---|---|---|
| NCC_ICIs (WES) | LUAD | TP53 | MUT/WT | 20 | 11 | 2.03 | 6.23 | 2.14 (0.51-9.03) | 0.3 | 28.9 | 32.9 | 1.49 (0.39-5.64) | 0.56 |
| NCC_ICIs (WES) | LUAD | KRAS | MUT/WT | 20 | 2 | 2.03 | 4.13 | 0.31 (0.04-2.58) | 0.28 | 28.9 | 32.9 | 0.51 (0.06-4.15) | 0.53 |
| NCC_ICIs (WES) | LUAD | STK11 | MUT/WT | 20 | 1 | 2.03 | 6.23 | 1.98 (0.22-17.85) | 0.54 | 28.9 | 32.9 | 1.71 (0.2-14.44) | 0.62 |
| NCC_ICIs (WES) | LUAD | KEAP1 | All WT | 20 | 0 | | | | | | | | |
| NCC_ICIs (WES) | LUAD | EGFR | MUT/WT | 20 | 10 | 10.38 | 4.13 | 0.81 (0.21-3.06) | 0.76 | 26.93 | 32.9 | 1.48 (0.39-5.71) | 0.57 |
| NCC_ICIs (WES) | LUAD | TP53#KRAS | MUT/WT | 20 | 1 | 2.03 | 6.23 | 1.98 (0.22-17.85) | 0.54 | 28.9 | 32.9 | 1.71 (0.2-14.44) | 0.62 |
| NCC_ICIs (WES) | LUAD | TP53#STK11 | MUT/WT | 20 | 1 | 2.03 | 6.23 | 1.98 (0.22-17.85) | 0.54 | 28.9 | 32.9 | 1.71 (0.2-14.44) | 0.62 |
| NCC_ICIs (WES) | LUAD | TP53#KEAP1 | All WT | 20 | 0 | | | | | | | | |
| NCC_ICIs (WES) | LUAD | TP53#EGFR | MUT/WT | 20 | 6 | 10.35 | 4.13 | 1.72 (0.42-6.95) | 0.45 | 26.93 | 32.9 | 2.19 (0.57-8.38) | 0.25 |
| NCC_ICIs (WES) | LUAD | KRAS#STK11 | MUT/WT | 20 | 1 | 2.03 | 6.23 | 1.98 (0.22-17.85) | 0.54 | 28.9 | 32.9 | 1.71 (0.2-14.44) | 0.62 |
| NCC_ICIs (WES) | LUAD | KRAS#KEAP1 | All WT | 20 | 0 | | | | | | | | |
| NCC_ICIs (WES) | LUAD | KRAS#EGFR | MUT/WT | 20 | 1 | NA | 2.03 | 0 (0-Inf) | 1 | | 28.9 | 0 (0-Inf) | 1 |
| NCC_ICIs (WES) | LUAD | STK11#KEAP1 | All WT | 20 | 0 | | | | | | | | |

| | | | | | | | | | |
|---|---|---|---|---|---|---|---|---|---|
| NCC_ICIs (WES) | LUAD | STK11#EGFR | All WT | 20 | 0 | | | | |
| NCC_ICIs (WES) | LUAD | KEAP1#EGFR | All WT | 20 | 0 | | | | |
| NCC_ICIs (WES) | LUAD | TP53#KRAS#STK11#KEAP1#EGFR | All WT | 20 | 0 | | | | |
| Wang_2019 (Panel) | LUAD | TP53 | MUT/WT | 28 | 14 | 2.73 | 3.23 | 1.04 (0.42-2.59) | 0.93 |
| Wang_2019 (Panel) | LUAD | KRAS | MUT/WT | 28 | 9 | | 2.77 | 0.29 (0.09-0.88) | 0.029 |
| Wang_2019 (Panel) | LUAD | STK11 | All WT | 28 | 0 | | | | |
| Wang_2019 (Panel) | LUAD | KEAP1 | MUT/WT | 28 | 1 | 2.8 | 3.13 | 1.66 (0.21-12.79) | 0.63 |
| Wang_2019 (Panel) | LUAD | EGFR | MUT/WT | 28 | 4 | 2.77 | 3.02 | 0.84 (0.19-3.64) | 0.81 |
| Wang_2019 (Panel) | LUAD | TP53#KRAS | MUT/WT | 28 | 6 | 2.9 | 2.8 | 0.45 (0.13-1.57) | 0.21 |
| Wang_2019 (Panel) | LUAD | TP53#STK11 | All WT | 28 | 0 | | | | |
| Wang_2019 (Panel) | LUAD | TP53#KEAP1 | MUT/WT | 28 | 1 | 2.8 | 3.13 | 1.66 (0.21-12.79) | 0.63 |
| Wang_2019 (Panel) | LUAD | TP53#EGFR | MUT/WT | 28 | 2 | NA | 2.85 | 0 (0-Inf) | 1 |
| Wang_2019 (Panel) | LUA | KRAS#STK11 | All WT | 28 | 0 | | | | |

| | | | | | |
|---|---|---|---|---|---|
| | D | | | | |
| Wang_2019 (Panel) | LUAD | KRAS#KEAP1 | All WT | 28 | 0 |
| Wang_2019 (Panel) | LUAD | KRAS#EGFR | All WT | 28 | 0 |
| Wang_2019 (Panel) | LUAD | STK11#KEAP1 | All WT | 28 | 0 |
| Wang_2019 (Panel) | LUAD | STK11#EGFR | All WT | 28 | 0 |
| Wang_2019 (Panel) | LUAD | KEAP1#EGFR | All WT | 28 | 0 |
| Wang_2019 (Panel) | LUAD | TP53#KRAS#STK11#KEAP1#EGFR | All WT | 28 | 0 |

[1]NoOfPts: number of all patients;
[2]NoOfMut: number of mutated patients;
[3]PFS_m_mut: median PFS of mutated patients;
[4]PFS_m_wt: median PFS of wide-type patients;
[5]PFS_HR95CI: HR and 95%CI of PFS;
[6]PFS_Cox_P: P value of PFS in Cox regression;
[7]OS_m_mut: median OS of mutated patients;
[8]OS_m_wt: median OS of wide-type patients;
[9]OS_HR95CI: HR and 95%CI of OS;
[10]OS_Cox_P: P value of OS in Cox regression;

**Table S3. The association of STK11 mutation with PFS or OS in each ICIs and non-ICI datasets.**

| Type | Cohort | Number of all samples | Number of mutated samples | PFS | | | | OS | | | |
|---|---|---|---|---|---|---|---|---|---|---|---|
| | | | | mPFS (mut[1]) | mPFS (wt[2]) | PFS HR (95%CI) | P value (PFS) | mOS (mut[1]) | mOS (wt[2]) | OS HR (95%CI) | P value (OS) |
| ICI-Cohort | Hellmann_2018 (WES) | 59 | 12 | 6.51 | 7.82 | 1.28 (0.58-2.79) | 0.54 | | | | |
| | Gandara_2018 (FM) | 301 | 32 | 1.41 | 2.79 | 1.54 (1.04-2.26) | 0.03 | 7.33 | 15.61 | 1.81 (1.19-2.74) | < 0.01 |
| | Rizvi_2015 (WES) | 29 | 5 | 9.5 | 12.6 | 3.14 (0.68-14.41) | 0.12 | | | | |
| | Rizvi_2018 (MSK-IMPACT) | 185 | 49 | 2.5 | 4 | 1.26 (0.89-1.78) | 0.20 | | | | |
| | Samstein_2018 (IMPACT) | 266 | 66 | | | | | 6 | 13 | 1.47 (1.05-2.06) | 0.03 |
| | Miao_2018 (WES) | 50 | 6 | 1.8 | 5.87 | 2.4 (0.92-6.27) | 0.06 | 4.4 | 15.73 | 3.48 (1.12-10.83) | 0.02 |
| | Bai_2020 (WES) | 50 | 1 | 4.23 | 2.1 | 0.84 (0.11-6.17) | 0.87 | | | | |
| | NCC_ICIs (WES) | 20 | 1 | 2.03 | 6.23 | 1.98 (0.22-17.85) | 0.53 | 28.9 | 32.9 | 1.71 (0.2-14.44) | 0.62 |
| | NCC | 202 | 13 | | | | | | | 1.94 (0.69-5.46) | 0.20 |

| | | | | | | | | | | |
|---|---|---|---|---|---|---|---|---|---|---|
| Non-ICI Cohort | Chen_2019 | 293 | 15 | | | | | 35.33 | 114.9 | 2.45 (1.18-5.09) | 0.01 |
| | TCGA_LUAD | 504 | 45 | 26.97 | 40.07 | 1.22 (0.76-1.95) | 0.41 | 39.8 | 49.97 | 1.23 (0.76-2.01) | 0.40 |

[1]mut: mutation; [2]wt: wide-type
