## Supplemental Tables for "IMPACT: a web server for exploring immunotherapeutic predictive and cancer prognostic biomarkers"

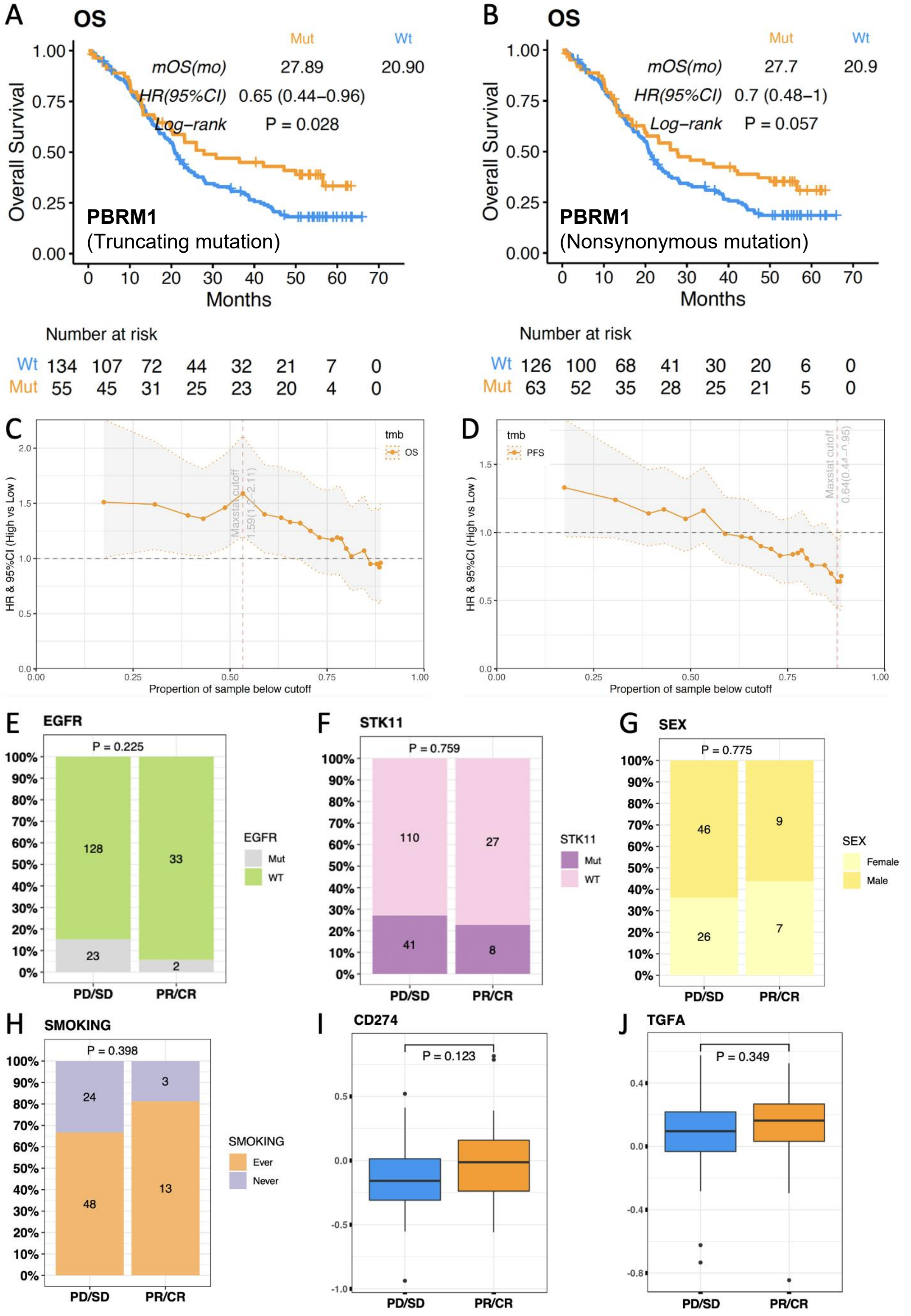


**Figure S1.** Examples of unique functions of IMPACT. **(A)** The association of *PBRM1* truncating mutation with OS in the kidney cancer dataset of Checkmate_025. **(B)** The association of *PBRM1* nonsynonymous mutation with OS in the kidney cancer dataset of Checkmate_025. **(C)** The hazard ratios of bTMB and OS across various cutoffs on Gandara_2018 dataset. **(D)** The hazard ratios of bTMB and PFS across various cutoffs on the Gandara_2018 dataset. **(E) - (F)** The bar plots of objective response rate against gene mutations on the LUAD dataset of Rizvi_2018. P values were calculated with the Chisq-test. **(G) - (H)** The boxplots of objective response rate against gene expression on the BLCA dataset of GSE176307. P values were calculated with the Mann Whitney U test. **(I) - (J)** The bar plots of objective response rate against clinical factors on the BLCA dataset of GSE176307. P values were calculated with the Chisq-test.

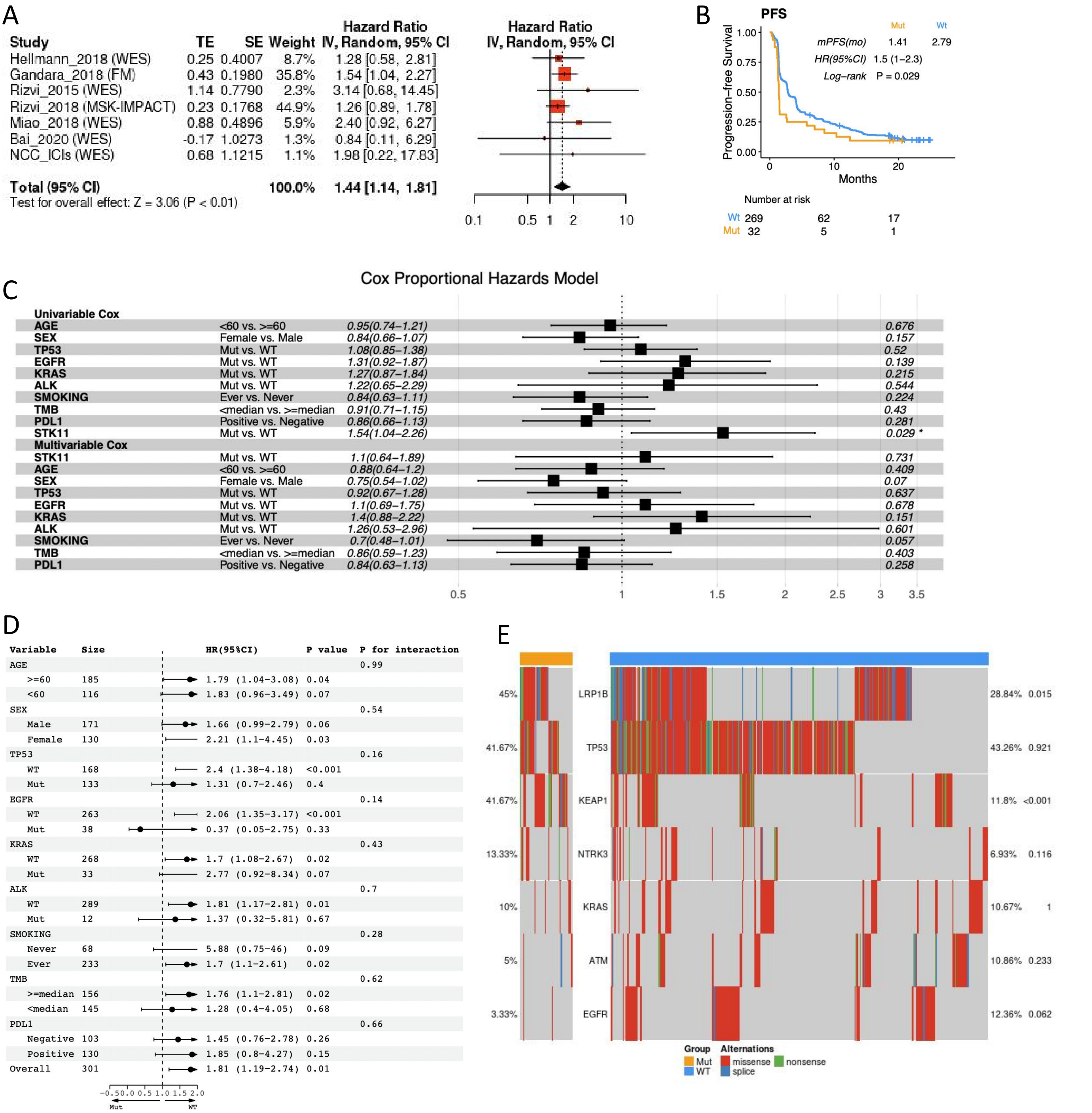


**Figure S2.** The exploration of *STK11* mutation as an ICI predictive biomarker. **(A)** Meta-analysis of *STK11* mutation and PFS across LUAD ICIs cohorts. **(B)** The Kaplan-Meier curve of *STK11* mutation and PFS on the Gandara_2018 dataset. **(C)** The univariable and multivariable cox regression of *STK11* mutation with PFS on the Gandara_2018 dataset. **(D)** Subgroup analysis of *STK11* mutation and OS on the Gandara_2018 dataset. **(E)** The correlation of *STK11* mutation with other genes mutations on the Gandara_2018 dataset.
